# Sensory adaptation in the barrel cortex during active sensation in the awake, behaving mouse

**DOI:** 10.1101/2022.07.07.499259

**Authors:** Andrea Colins Rodriguez, Michaela S. E. Loft, Ingo Schiessl, Miguel Maravall, Rasmus Petersen

## Abstract

Sensory adaptation (SA) is a prominent aspect of how neurons in anaesthetised animals respond to sensory signals, ubiquitous across species and modalities. However, SA depends on the activation state of the brain and it has been doubted whether SA is expressed in behaving animals. Here, we addressed this question by training mice to detect an object using their whiskers and recording neuronal activity from barrel cortex whilst simultaneously imaging the whiskers in 3D. We found that neuronal responses decreased during the course of whisker-object touch sequences but that this was due to two factors. First, a motor effect, whereby, during a sequence of touches, later touches were mechanically weaker than early ones. Second, a sensory encoding effect, whereby neuronal tuning to touch became progressively less sensitive during the course of a touch sequence. The sensory encoding effect was whisker-specific. These results show that SA does occur during active whisker sensing and suggest that SA is fundamental to sensation during natural behaviour.

## Introduction

Sensory Adaptation (SA), first demonstrated in single neurons by Adrian & Zotterman (1926), is exemplified by the phenomenon that the response of neurons to a sensory stimulus of fixed strength (e.g., whisker deflection or light flash) progressively decreases during the course of a sequence of such stimuli. In general, SA refers to phenomena whereby neuronal responses are dynamic – depending not only on the immediate stimulus but also on the temporal context in which they occur (Ahissar et al., 2000; Chung et al., 2002; Díaz-Quesada & Maravall, 2008; Lampl & Katz, 2017; Laughlin, 1989; Maravall, 2013; Maravall et al., 2007, 2013; Petersen et al., 2009; Ulanovsky et al., 2004; Wang et al., 2010; Weber et al., 2019; Wright et al., 2021).

SA has been thoroughly studied under anaesthetised conditions and, to a lesser extent, in awake animals with immobilised sense organs (Lampl & Katz, 2017; Weber et al., 2019). In the ‘passive stimulation’ paradigm, stimuli are applied to the immobilised animal – the animal is the passive recipient of stimulation. This paradigm is experimentally convenient for studying SA, since the experimenter has a high degree of control over delivery of sensory stimuli. However, the experimental conditions differ from those of behaving animals in important respects.

Behaving animals acquire sensory information by making exploratory movements of their sense organs (Active Sensing, AS) (Gibson, 1962; Yarbus, 1967). For example, primates explore scenes by eye movements and rats/mice by whisker movements. The value of AS is that it allows animals to make actions that sample sensory information most useful to the animal’s current goals and thus to attain those goals more efficiently than would be possible via passive sensing (Prescott et al., 2011). Human eye movements can approach the efficiency of an ideal Bayesian planner (Yang et al., 2016) and rats/mice use whisker movements to turn challenging tactile discrimination tasks into more straightforward detection tasks (Campagner et al., 2018; O’Connor et al., 2010). However, it is technically challenging to study SA during AS. In contrast to the passive stimulation paradigm, where SA can be straightforwardly assayed by comparing the response of neurons to a sequence of stimuli of fixed strength, during AS, animals continually move their sense organs and thereby change the sensory input. This presents a significant challenge for the investigation of SA: a change in neuronal response during the course of a sequence of whisker touches might not be due to SA but rather to a change in touch strength. Whether SA is expressed in the actively sensing animal engaged in a goal-driven task is an open question.

Here we address this issue by studying the mouse whisker system and taking advantage of recent advances in methods for quantifying whisker-dependent behaviour (Campagner et al., 2018; Clack et al., 2012; Petersen et al., 2020). The whisker system exhibits AS in a tractable form – rats/mice explore objects by probing them with back-and-forth movements of their whiskers (“whisking”). There is consistent evidence for SA in the whisker-related zone of primary somatosensory cortex of the anaesthetised rat/mouse (Chung et al., 2002; Ganmor et al., 2010; Katz et al., 2006; Khatri et al., 2004, 2009; Maravall et al., 2007, 2013; Musall et al., 2017; Simons, 1985) and this SA is specific to individual whiskers (Castro-Alamancos, 2004; Crochet et al., 2011; Crochet & Petersen, 2006; Erchova et al., 2002; Fanselow & Nicolelis, 1999; Katz et al., 2006; Yamashita et al., 2013). However, findings from awake animals have been inconsistent, with some studies reporting SA to be substantial and others reporting it to be weak/absent (Castro-Alamancos, 2004; Crochet et al., 2011; Crochet & Petersen, 2006; Fanselow & Nicolelis, 1999; Hentschke et al., 2006; Musall et al., 2014; Wright et al., 2021; Yamashita et al., 2013).

To test whether SA occurs in the behaving, actively sensing mouse, we trained mice to perform a whisker-guided object detection task, and then recorded neural activity from barrel cortex whilst, simultaneously, we imaged their whiskers in 3D. We found that cortical responses to whisker-object touch do depend on prior touches. This effect originates not only from a direct effect on the responsiveness of sensory neurons (SA) but also from an indirect effect whereby touch triggers a change in motor action which in turn affects the strength of future touches.

## Results

### Variability of whisking behaviour

Our first aim was to establish a behavioural paradigm to test whether Sensory Adaptation (SA) occurs during active touch. Head-fixed mice can discriminate the location of a pole using their whiskers and use active touch to do so (Campagner et al., 2018; O’Connor et al., 2010). As the mouse repeatedly moves its whiskers back and forth against the pole, this results in touch sequences with inter-touch intervals of up to about 100 ms (Campagner et al., 2018; O’Connor et al., 2010). It is well-established that sequences of touches with such intervals, delivered passively to anaesthetised mice, produce strong SA in barrel cortex (Chung et al., 2002; Simons, 1985). We therefore reasoned that, if SA is expressed in the awake, actively sensing animal, then it should be engaged by such behaviour. To test this end, mice (N=3) were trained to solve a Go-NoGo pole detection task (Figure 1A) with a single row of whiskers (C1, C2 and C3). On ‘Go’ trials, the pole was moved within reach of the whiskers: if the mouse licked the lickport (hit), the mouse was rewarded with a water droplet; if not (miss), it was penalised with a time-out. On ‘NoGo’ trials, the pole was placed out of reach of the whiskers: if the mouse withheld licking (correct rejection), the task proceeded immediately to the next trial; if not (false alarm) the mouse was penalised with a time-out and sound. As previously reported (Petersen et al., 2020), mice learned to perform the task accurately (81±17%, mean ± SD over mice), performing 135±22 trials per session.

**Figure 1:**
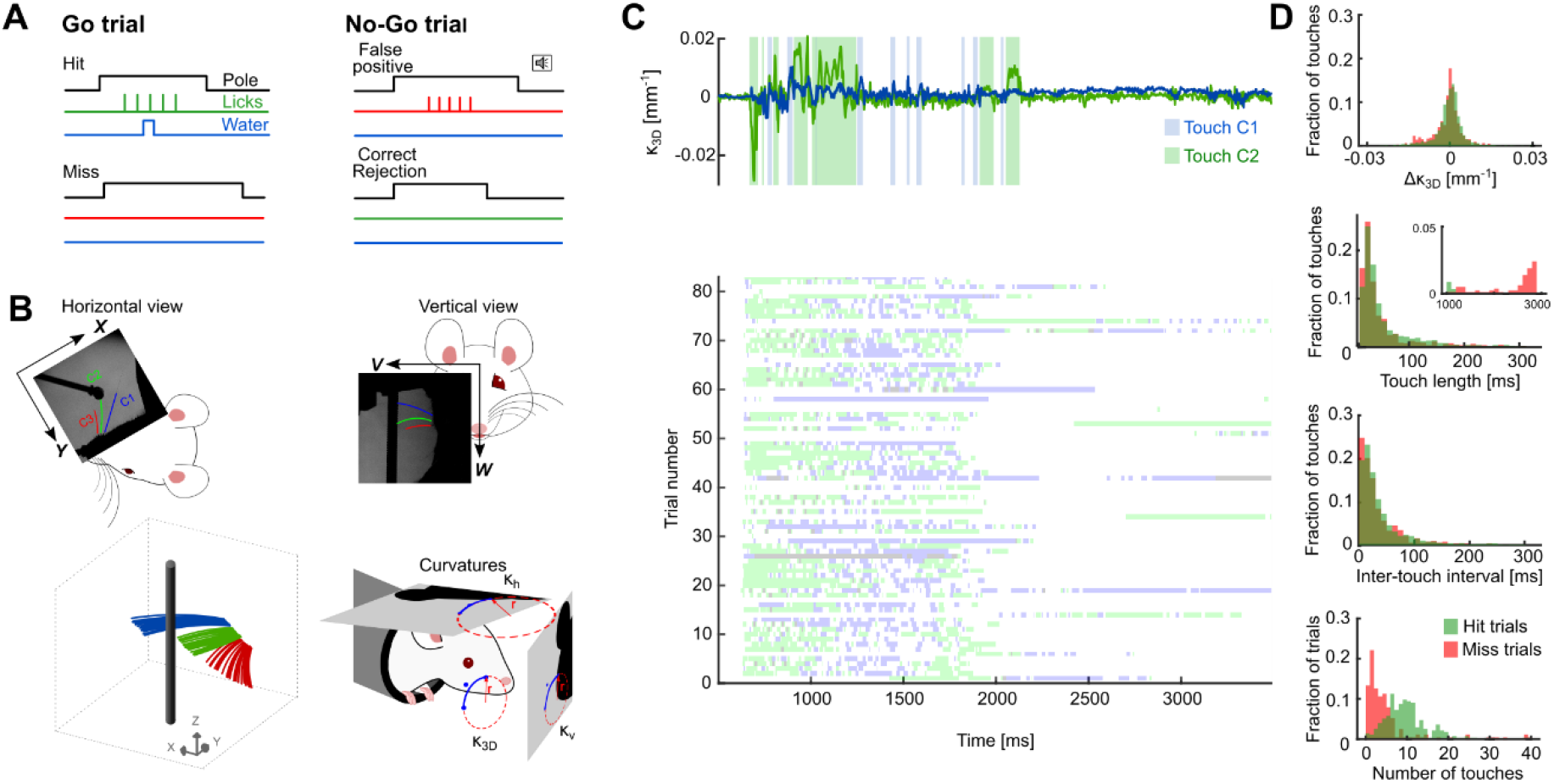
Variability of whisker-pole touches during the pole detection task. **A)** Behavioural task schematic. Mice were trained to report the pole’s presence within whisker reach (Go trials) by licking a port. On NoGo trials, the pole was moved to a position out of whisker reach and mice should withhold licks. False alarms were penalised with a tone, while Misses were penalised with a time-out period. **B)** Whisker imaging. Whiskers (C1, C2 and C3) were imaged with two cameras during the behavioural task (top panels). Whiskers were tracked in 3D and 3D curvature (*κ*_3*D*_) was then calculated (bottom panels). **C)** Top: Whisker-pole touches and *k*_3*d*_ from an example Go trial. Bottom: Touches for an example session (in this session only C1 and C2 whiskers touched the pole). **D)** Behavioural variability of touches (hit trials green; miss trials red): Strength of touch (Δ*k*_3*D*_), touch-length (inset shows fraction of touches with contact periods >1000 ms), inter-touch interval and number of touches per trial.

In order to measure whisking behaviour in 3D, we imaged the whiskers at 1000 frames/s with two video cameras (Figure 1B). Using a 3D whisker tracking algorithm (Petersen et al., 2020), both 3D whisker shape and 3D whisker position were extracted from the imaging data (total of 1.8M frames). From whisker shape, the 3D curvature (*κ*_3*D*_) at the base of each whisker was estimated for each frame (Figure 1B). We estimated the strength of whisker-pole touch in each frame as the change in curvature at the whisker base relative to its intrinsic curvature (Δ*k*_3*D*_).

As desired, each trial consisted of a sequence of one or more ‘touches’ (a contiguous sequence of frames in which the whisker was in continuous contact with the pole) (Figure 1C). In contrast to the regular stimulus sequences usually used in sensory physiology, we found that the temporal pattern of touches varied substantially across trials. This was manifest in the number of touches per trial, the interval between touch onsets, the duration of touch, the strength of touch (Δ*k*_3*D*_) and the direction of touch (sign of curvature change) (Figure 1D). Touches on miss trials were similar in duration, interval and strength to those on hit trials but the number of touches per trial was less (10.5 ± 5.7 per trial on hit trials, 5 ± 7.7 on miss trials; Figure 1D). This observation raises the question of whether, in the behaving animal, SA to repeated touches is a strong enough effect to be significant above the ‘noise’ of behavioural touch variability or whether it might be a phenomenon only observable when touch variability is unusually low, as under passive whisker stimulation.

### Response attenuation during a touch sequence

To determine if SA occurs during active touch, we made extracellular recordings of neural activity (single-unit, N=33 obtained from 6 sessions) from barrel cortex columns C1-C3 (corresponding to the spared whiskers) whilst mice performed the pole detection task. Units showed a clear response to touch (Figure 2A and Figure 2C). For each trial, we classified touches according to whether they were first, second, third, etc on that trial. We compared responses across touches and between hit and miss trials by 2-way ANOVA (main effects of first vs later touch and hit vs miss trial). We found that the first touch evoked a substantially stronger response than later touches (p<10e-6), and that there was a progressive attenuation in response (firing rate) during the course of a series of touches (Fig 2A-C). Responses were similar on error (miss) trials: (ratio of firing rate on miss vs hit trials 1.06; p=0.06; Figure 2D), indicating that this response attenuation is independent of central processing of trial outcomes. First touch responses were not only stronger than those of later touches but also more discriminable from spontaneous activity (quantified by area under ROC curve; 2-way ANOVA; main effect of first vs later touches, p<10^-6; main effect of hit vs miss, p=0.53; Figure 2E). We refer to this phenomenon as ‘response attenuation’. As detailed below, it is important to distinguish response attenuation from SA, which is just one of several factors that could potentially account for it.

**Figure 2.**
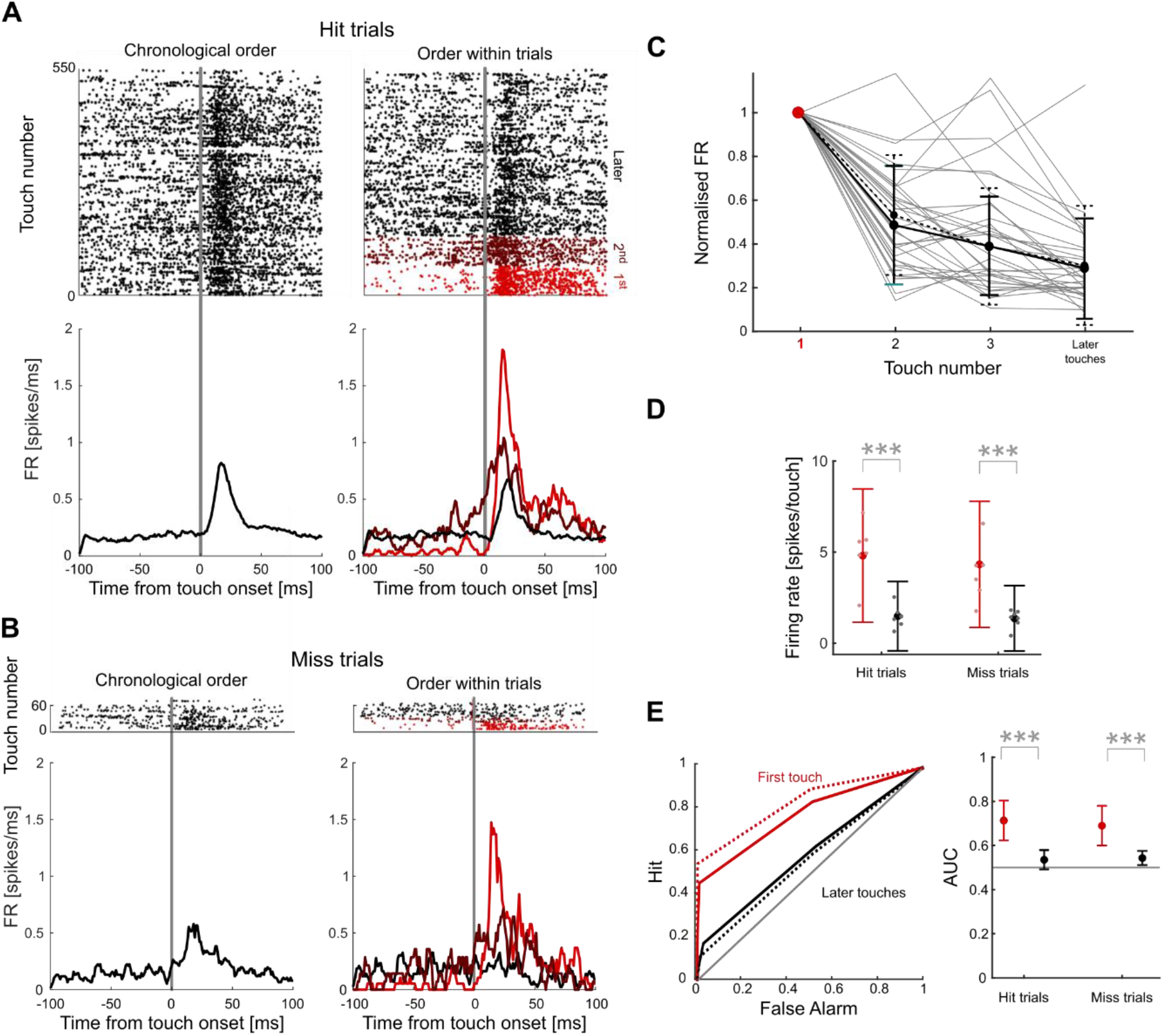
Response attenuation in barrel cortex during touch sequences. **A)** Raster plots and PSTHs of the neural population recorded during one session divided into hit trials. Left panels show data sorted by chronological order of touches, while right panels show the same data sorted by touch order within trials (first touch, second touch etc). PSTHs were smoothed (5 ms boxcar) for display only. **B)** Same as A for miss trials. **C)** Normalised responses of all units to touch ordered by their occurrence within trials (grey lines). Firing rate was normalised by responses to the first touch. The black solid line represents mean and SD across neurons in hit trials and the dashed line represents mean and SD across neurons in miss trials. **D)** Comparison of responses to touches (spike count within 30 ms after onset) during hit and miss trials, across the sample of units. Small dots indicate averages within sessions, large dots show the mean across all units and error bars indicate SD across all units. **E)** Left: Receiver Operating Characteristic (ROC) curve of touch detection for an example unit. Diagonal line represents chance performance (area under ROC curve is 0.5). Solid lines represent ROC for hit trials; dashed lines for miss trials. Right: Area under ROC curve for first and later touches of hit and miss trials across all units. Grey line represents chance performance. For all panels: * indicates p-value<0.05, ** indicates<0.005, ***p-values<0.0005.

### Motor adaptation

Although these response attenuation data are consistent with the hypothesis that SA is elicited by active touch during our task, there is a potential alternative explanation. Since the touches in these touch sequences are far from identical (Figure 1), the decrease in firing rate from first to later touches might potentially be due to physical differences in stimulus strength between first and later touches. To test this possibility, we used 3D whisker tracking to measure both how far, and how fast, the whiskers were displaced during touches, and also how much the whiskers bent during touches (Figure 3AB). An advantage of our 3D whisker imaging method is that the effect of bending on whisker shape can be teased apart from that of 2D image projection artefacts (Knutsen et al., 2008; Petersen et al., 2020).

**Figure 3:**
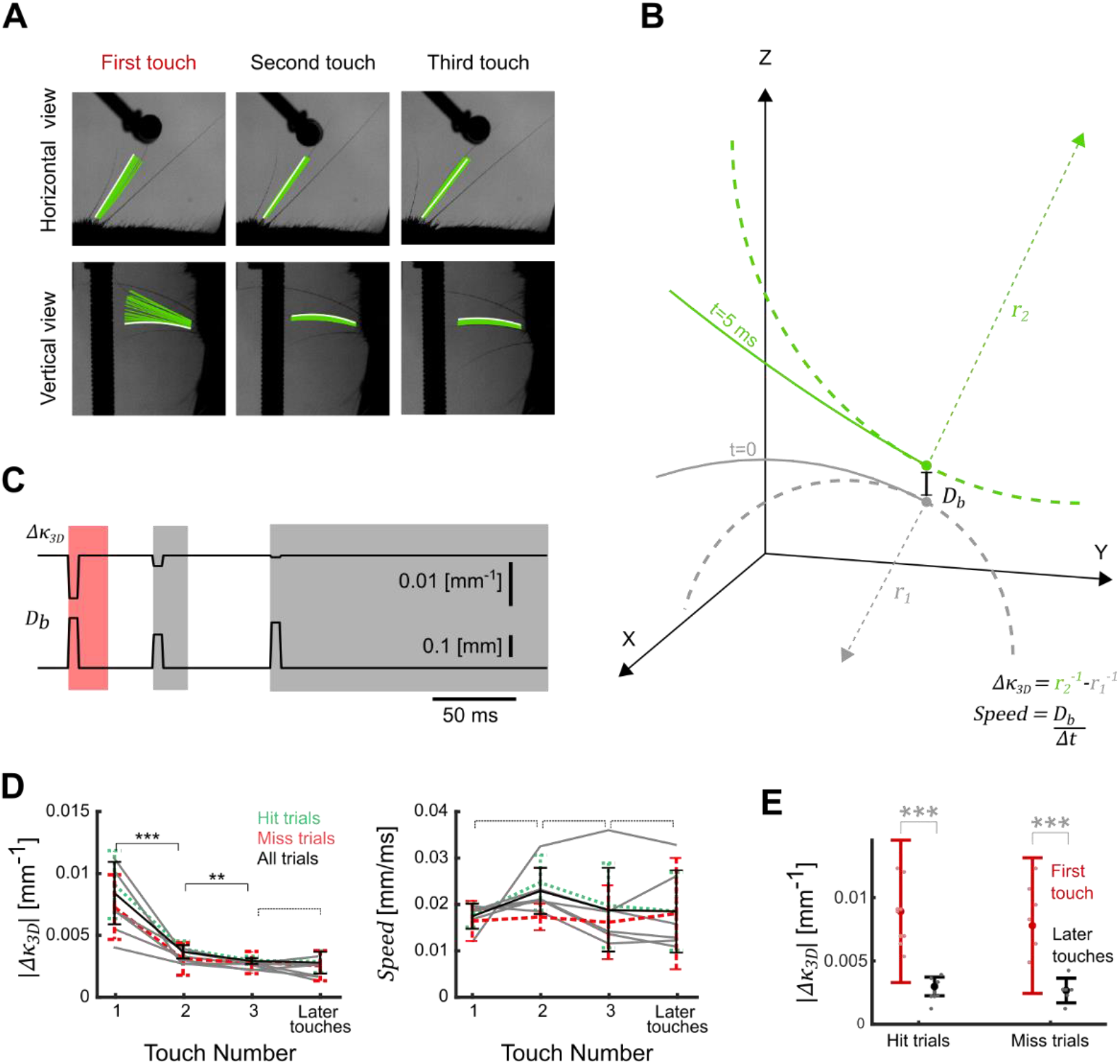
Strength of whisker touch decreases during a touch sequence. **A)** Single-trial example showing whisker tracking solutions (Bezier curves) projected onto both horizontal and vertical imaging planes. Tracking solution at touch onset is white, whilst solutions for the subsequent 20 ms are green. **B)** Extraction of whisker shape/displacement parameters from the tracking solutions. Bezier curves describe a whisker at touch onset (*t*=0, grey solid curve; circle marks whisker base) and 5 ms later (*t*=5, green solid curve). Whisker displacement (*D*_*b*_) is the distance between whisker base at *t*=0 and that at *t*=5, and is proportional to whisker speed. Curvature of the whisker at its base is derived from the length of the radius (*r*_1_; dashed grey line) of the circle (dashed grey curve) whose curvature matches that of the whisker at the base. Dashed green line/curve, with associated radius *r*_*2*_, show corresponding data for *t*=5. Touch magnitude (Δ*k*_3*D*_) is the difference in the reciprocals of *r*_1_ and *r*_*2*_. *X, Y* and *Z* axes correspond to the coordinates defined in Figure 1B). **C)** Δ*k*_3*D*_ and *D*_*b*_(mean of their values in *t*=[0,5]) for each of the three touches shown in panel A. Shaded areas indicate touches. **D)** Left: |Δ*k*_3*D*_| measured as in panel B as a function of touch number for each session (grey lines), averaged across all trials (black), hit trials (green) and miss trials (red); error bars denote SD across trials. Right: Corresponding data for whisker base speed (*D*_*b*_/*dt*). E) Comparison of |Δ*k*_3*D*_| for correct and miss trials. Pink dots indicate averages within a session, red dots and error bars are the average and standard deviation over all trials (from all sessions) respectively. For all panels: * indicates p-value<0.05, ** indicates p-values<0.005, ***p-values<0.0005. Dashed brackets indicate no significant difference.

We compared whisker speed and touch strength (Δ*k*_3*D*_), computed at the whisker base, of first touches to later touches. We tested the effect of first vs later touch and of hit vs miss trial on these variables by 2-way ANOVA. We found that whisker speed was similar but that there was a substantial decrease in Δ*k*_3*D*_ (main effect of first vs later touch on Δ*k*_3*D*_, p<1e-6; on *D*_*b*_, p=0.9; Figure 3C). These data indicate that active touch on this task involves pronounced ‘motor adaptation’, whereby animals make progressively weaker pole contact during the course of a trial. The evolution of Δ*k*_3*D*_ and *D*_*b*_was similar on hit and miss trials (main effect of hit vs miss, p>0.8 for Δ*k*_3*D*_, p=0.27 for *D*_*b*_; Figure 3D), indicating that, as with the response attenuation reported above, the motor adaptation effect is independent of central processing of trial outcome. Since whisker bending is the best single predictor of the response of whisker mechanoreceptors to touch (Bush et al., 2016; Campagner et al., 2016; Severson et al., 2017; Tonomura et al., 2015), these data imply a decrease in strength of afferent drive to barrel cortex during the course of a trial. This suggests that the attenuation of cortical response reported above (Figure 2) need not be an entirely due to SA but could, either entirely or in part, be an indirect effect of motor adaptation.

### Sensory Adaptation

To determine whether SA contributes to response attenuation, we developed a method to test for potential changes in SA during the course of a touch sequence. The natural variability in touch reported in Figure 1, allowed us to estimate tuning curves to touch strength. Consistent with previous studies (Hires et al., 2015; Peron et al., 2015; Petreanu et al., 2012; Sofroniew et al., 2015), we found that barrel cortical units were typically tuned to whisker bending: the greater the bending (Δ*k*_3*D*_), the greater the firing rate (Figure 4A). If SA is indeed expressed, the prediction is that tuning curves (firing rate as a function of Δ*k*_3*D*_) should shift downwards (units become less sensitive to touch) during the course of a touch sequence. We used linear regression to estimate tuning curves, since we found linearity to be a good approximation within the range of Δ*k*_3*D*_ that we were considering (Figure 4AB).

**Figure 4:**
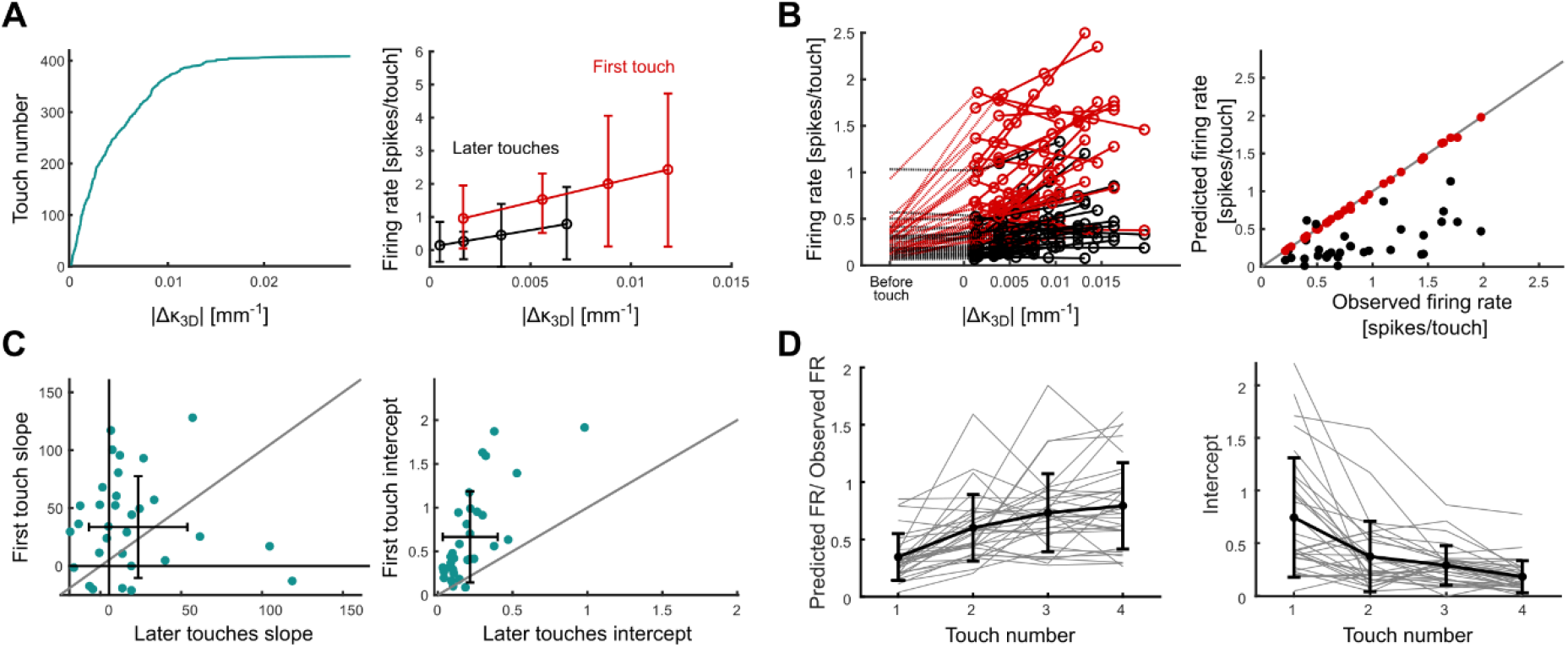
Sensory Adaptation. **A)** Left: sorted values of Δ*k*_3*D*_ in an example session. Right: Tuning curves for first and later touches for an example unit recorded during session shown in A. **B)** Left: Tuning curves for first and later touches for all selected units. Responses in the 30 ms interval before touch are shown for comparison. Right: Prediction of response to first touch using tuning curves estimated for later touch (black circles) and first touch respectively (red circles). **C)** Comparison of tuning curves parameters for first and later touches. Left panel shows slope and right panel shows intercept of tuning curves for all selected units. **D)** Left: Ratio between predicted firing rate by later touches tuning curve (*N*_*touch*_>4) and mean observed firing rate for each touch. Grey lines represent mean for each unit and black line represents average across units. Right: Intercept of tuning curves for each touch. Grey lines represent mean for each unit and black line represents average across units.

Figure 4A (right) shows results for an example unit. As predicted by the SA hypothesis, there was a downwards shift in tuning curves from first to later touches. This was a typical result (Figure 4B). To quantify the extent of this adaptation, we used the tuning curves to estimate and compare the responses of units to first vs later touches, keeping touch strength fixed. Specifically, we compared the actual response of each unit to first touch (red dots in Fig 4B right) to that predicted by feeding the touch strength (Δ*k*_3*D*_) of the first touch through the tuning curve estimated for later touches (black dots in Fig 4B right). For 85% of units, the responses to first touch predicted in this way were significantly lower than the actual responses (*t*-test, mean p-value=0.0023) - on average, 67% lower. Repeating this analysis for tuning curves estimated for\ *N*_*touch*_>4 in turn showed a gradual convergence to an adapted state (Figure 4D). Both the slope and intercept of the tuning function for later touch decreased significantly compared to those for first touch (Figure 4C), with the strongest effect observed for the intercept: 93% of units had higher intercept for first touch; 63% had higher slope for first touch. The intercept decreased gradually over the course of later touches (Figure 4D right). These results indicate that a significant part of the response attenuation from first to later touches observed in Figure 2 can be explained by a change in neuronal sensitivity. Thus, response attenuation is not simply a consequence of motor adaptation – there is also SA.

To estimate how much of the overall response attenuation in Figure 2 is due to SA, for each simultaneously recorded population of units, we first determined the overall response attenuation from first to later touch due to the combined effects of motor adaptation and SA (calculated from Figure 2B). Next, we estimated how much of this decrement could be attributed to the effect of tuning curves alone, keeping touch strength constant (Methods). We found that, on average across the sessions, 73% of the overall attenuation in firing rate is attributable to SA.

In sum, these results indicate that both Sensory Adaptation and Motor Adaptation contribute to the attenuation in firing rate observed during a sequence of whisker-pole touches, with Sensory Adaptation playing the major role.

### Stimulus-Specific Adaptation

A hallmark of sensory adaptation in studies of anaesthetised animals is that adaptation can be specific to individual whiskers (Katz et al., 2006; Musall et al., 2017). That is, stimulation of a given whisker attenuates responses to subsequent stimulation only of that whisker and not to that of other whiskers (whisker-specific adaptation, WSA). The hypothesis that SA is operative in the actively sensing, behaving animal suggests, therefore, that the changes in tuning curve that we reported above should be whisker-specific. To probe for WSA, we selected all trials in a session in which at least the first two touches were produced by the same whisker *W*_*ad*_ (C1 or C2) and at least one later touch (3rd or later) was made by a different whisker *W*_*test*_. Touch produced by whisker *W*_*ad*_ was considered as the adapting stimulus; touch produced by whisker *W*_*test*_ as the test stimulus (Figure 5A). Simultaneous touches were excluded. WSA predicts that the test stimulus should evoke an unadapted response.

**Figure 5:**
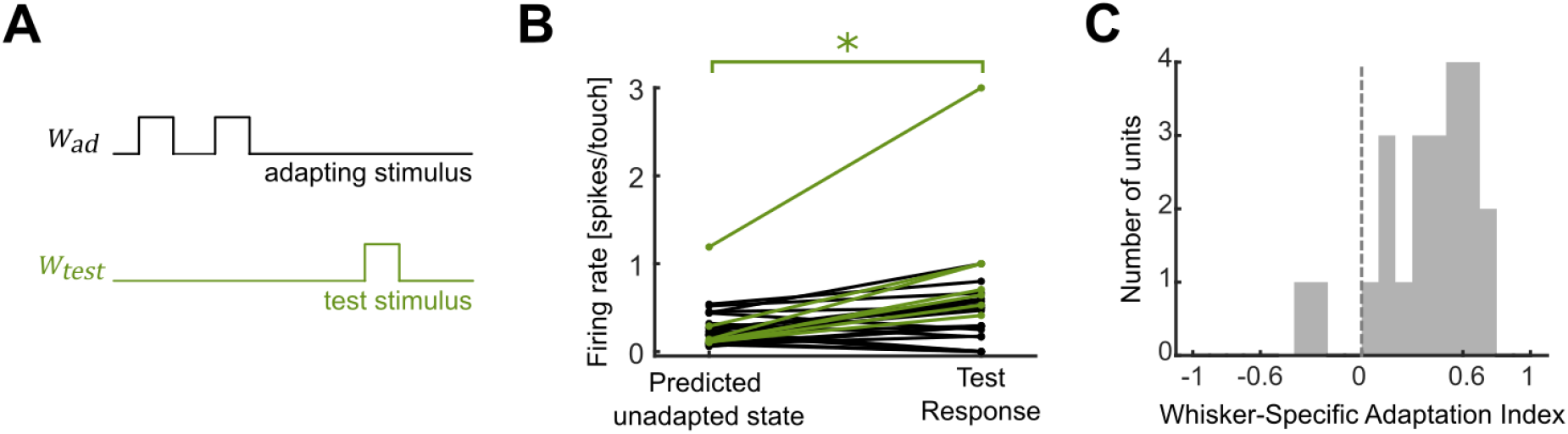
Whisker-specific Adaptation. A) Schematic of the analysis. If adaptation is whisker-specific, the response to the test stimulus should be greater than that to the same whisker in the adapted state (‘adapted response’). B) Comparison of response to the test stimulus with that predicted in the adapted state (see main text). Units whose responses were significantly higher than predicted in the adapted state highlighted in green. C) Histogram of SSA index (SI) values across the population.

To test this, we compared the responses of units to whisker *W*_*test*_ touches satisfying the definition of ‘test stimuli’ (*FR*_*test*_) (Fig. 5A) to whisker *W*_*test*_ in the adapted state (that is, responses to the third or later touch of that whisker within a trial). To factor out potential differences in touch strength, we measured tuning curves of units to |Δ*k*_3*D*_| of that whisker in the adapted state, and used these tuning curves to predict responses to touches of identical |Δ*k*_3*D*_| to those measured for the test stimuli (*FR*_*predicted*_). WSA predicts that *FR*_*test*_ should be greater than *FR*_*predicted*_ and that the following WSA index *I* should be positive:

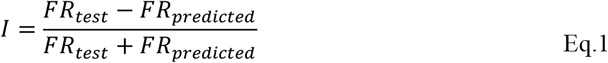

Alternatively, if adaptation spreads across whiskers, then *I* should be zero. We found that units had significantly higher response to the test stimulus compared to that predicted in the adapted state (Fig. 5B, *t*-test, p<0.05), that 91% of units had positive *I* (Fig. 5C) and that, across the population, *I* was significantly positive (t-test; p<2e-6). These results suggest that adaptation in our awake, actively sensing paradigm did exhibit WSA.

## Discussion

Our aim was to investigate the extent to which SA is expressed under conditions of active sensation, where the sensory input to the brain is determined by the animal’s own motor actions. Our main finding was that the response of neurons in barrel cortex decreased whilst a mouse repeatedly touched an object, and that this was due to two distinct factors: first, a motor adaptation effect, whereby mice tuned their whisking movements so that later touches were mechanically weaker than early ones; second, a sensory adaptation effect, whereby neurons became progressively less sensitive to touch. Thus, under conditions where animals are actively sensing the environment, the activity in cortical circuits reflects a delicate interplay between neuronal and behavioural dynamics.

Previous research on SA used ‘passive sensation’ paradigms, where the animal – usually anaesthetised - is a passive recipient of stimulation applied by the experimenter. The advantage of passive sensation is that the experimenter can deliver a train of identical stimuli and test for SA by comparing responses to them. Although it has long been recognised as important to test for SA in actively sensing, behaving animals, the challenge has been that awake animals move their sense organs and this makes it problematic to compare responses across repeated stimuli. Our approach was to train mice to perform a task (pole detection) that naturally elicits a stimulus sequence of whisker-pole touches. By tracking the whiskers, we showed that the resulting whisker-object touches are markedly variable in their temporal pattern, duration and strength. To disentangle the effects of this variance on neuronal activity in barrel cortex from that of SA, we used 3D whisker tracking to detect whisker-pole touches and to estimate their strength (Birdwell et al., 2007; Pammer et al., 2013; Petersen et al., 2020). This allowed us to measure how neurons are tuned to touch strength (the primary mechanical variable encoded by neurons in the ascending whisker pathway (Bush et al., 2016; Campagner et al., 2016; Hires et al., 2015; Peron et al., 2015; Petreanu et al., 2012; Severson et al., 2017; Sofroniew et al., 2015; Tonomura et al., 2015) and how this tuning adapts over the course of a touch sequence.

Previous work on SA in the anaesthetised brain or brain slice has found that cortical neurons strongly adapt when stimulated by trains of sensory input with a stimulus rate of a few Hz or more (Ahissar et al., 2000; Chung et al., 2002; Simons, 1985). However, of the few studies that have been conducted in awake animals, some report that SA is weak (Castro-Alamancos, 2004; Crochet et al., 2011; Crochet & Petersen, 2006) others that it is substantial (Musall et al., 2014; Ollerenshaw et al., 2014; Wright et al., 2021). The fact that SA can depend substantially on experimental conditions makes it important to study SA under conditions representative of natural behaviour. In particular, since the whisker system is an active sensory system, it is important to study SA whilst animals actively control their sensory apparatus to solve behavioural tasks, and to do so in a manner that controls for differences in stimulus strength. Our results indicate that SA is indeed expressed during the course of active whisker exploratory touch sequences. Controlling for differences in mechanical touch strength revealed that neurons were more sensitive to the first touch in a trial than to later touches. Thus, under physiologically representative conditions of active whisker exploration, adaptation significantly affects neuronal responses and tuning.

There are multiple mechanisms which contribute to SA, and SA is known to depend on a host of parameters - stimulus frequency, stimulus magnitude and brain state (Castro-Alamancos, 2004; Katz et al., 2012; Lampl & Katz, 2017; Wang et al., 2010). This suggests that the mechanisms expressed in any particular experiment are likely to depend on its precise conditions. The first mechanism to be clearly established was synaptic depression in thalamocortical synapses. For SA in the anaesthetised brain, this effect is prominent (Castro-Alamancos & Oldford, 2002; Chung et al., 2002). On the basis that thalamic activity is higher in activated vs quiescent brain states, it was argued that thalamocortical synapses are chronically depressed in the awake brain and, consequently, that there is little scope for thalamocortical synaptic depression to produce SA in the activated brain (Castro-Alamancos, 2004). However, subsequent research has found that mechanisms involving excitatory-inhibitory balance, subcortical adaptation and synchrony-sensitivity also contribute to SA (Chung et al., 2002; Gabernet et al., 2005; Lampl & Katz, 2017; Wang et al., 2010). Indeed, a recent study of awake mice, reported that, although SA could not be explained by synaptic depression of thalamocortical synapses, it could be explained by non-linear amplification of thalamic SA, due to the sensitivity of layer 4 cortical neurons to synchrony in thalamic activity (Wright et al., 2021). Taken together, these studies suggest that mechanisms different to synaptic depression gain in relative prominence under awake, behaving conditions.

Because active sensation occurs during arousal and exploration, the mechanisms whose dynamics underlie SA during active sensing must remain dynamic in the aroused state. As in other studies, we found that SA exhibited whisker-specificity, consistent with the notion that adaptation is caused by repeated activation of the specific pathways linking a follicle to the responding neurons (Lampl & Katz, 2017)

Active sensation is the general principle that animals use motor actions to acquire sensory information and that they select those actions expected to yield the sensory information most useful to their current goals (Yang et al., 2016). We found that the strength of whisker-pole contact decreased systematically during the course of a trial (motor adaptation). This is reminiscent of the ‘minimal impingement’ strategy whereby whisking amplitude decreases upon contact with an object (Mitchinson & Prescott, 2007). It is also consistent with extensive evidence for active sensing in animals with mobile whiskers, including rats, mice and aquatic mammals (Arkley et al., 2014; Campagner et al., 2019; Grant et al., 2013; Mitchinson et al., 2007; O’Connor et al., 2010; Sofroniew et al., 2014; Towal & Hartmann, 2006). Our finding that motor adaptation is similar on correct and incorrect trials suggests that its principal mechanism is insensitive to top-down signals from the brain’s decision-making centres and may involve anatomical loops relatively close to the periphery of the sensorimotor system.

## Methods

All experimental protocols described in here were approved by both United Kingdom Home Office national authorities and institutional ethical review.

### Behavioural apparatus

The whiskers were imaged similarly to (Petersen et al., 2020) using two high-speed cameras (LTR2, Mikrotron, Unterschleissheim, Germany; 1000 frames/s, 0.2 ms exposure time) with telecentric lenses (55-349, Edmunds Optics, Barrington, NJ). In relation to standard anatomical planes for the skull, one camera (480 × 480 pixels) imaged the horizonal plane. The second camera imaged a plane 25^°^ off the sagittal plane and 10^°^ off the horizontal plane; chosen to minimise occlusion between the cheek of the mouse and the whiskers. Mice were head-fixed and positioned inside a perspex tube (inner diameter 32 mm). The whiskers were free to move at all times. Whiskers were illuminated from below using an infrared LED array (940 nm, LED 940-66-60, Roithner, Vienna, Austria) via a diffused condenser lens.

### Surgical procedure and mice training

Mice (C57; males; N=3) were first habituated to the apparatus. To implant the titanium head-bar, mice were anaesthetised with isoflurane (1.5-2.5% isoflurane by volume in *0*_2_) while body temperature was maintained at 37°C using a homeothermic heating system. The mouse’s skull was exposed and the titanium head-bar was attached with dental acrylic. After surgery, mice were left to recover for at least 5 days before starting water restriction (1.5 ml water/day). Training began 7-10 days after the start of water restriction. Water was given as a reward during the training. Mice were trained until they reached a performance higher than 70%. Subsequently, a craniotomy was performed and sealed with silicone elastomer. Mice were left to recover for 2-3 days until performance had recovered.

### Behavioural task

Mice were trained on the pole detection task detailed in Petersen et al (2020). Briefly, mice were trained and imaged in a dark, sound-proofed enclosure. A head-fixed mouse was placed inside a perspex tube, from which its head emerged at one end. The stimulus object was a 0.2 mm diameter, vertical carbon fibre pole which could be translated in the horizontal plane by stepper motors. To allow vertical movement of the pole into and out of range of the whiskers, the apparatus was mounted on a pneumatic linear slide, powered by compressed air. The apparatus was controlled from MATLAB via a real-time processor. Mouse response was monitored by a lick port located anterior to the mouth. Licks were detected as described in (Campagner et al., 2019). Each lick port consisted of a metal tube connected to a water reservoir via a computer-controlled solenoid valve. Head-fixed mice were trained to detect the presence of the pole using their whiskers, using behavioural procedures similar to (Campagner et al., 2019). On each trial, the pole was presented either within reach of the whiskers (‘Go trial’) or out of reach (‘NoGo trial’). At the start of each trial, the computer triggered the pole to move up (travel time ∼100 ms). The pole stayed up for 1 s, before moving down. On Go trials, the correct response was for the mouse to lick a lick port. Correct responses were rewarded by a drop of water (∼10 μl). Incorrect responses on Go trials (not licking) were punished by timeout (3–5 s). On NoGo trials, the correct response was to refrain from licking and incorrect responses (licking) were punished by timeout and tone (frequency 12 kHz).

### Electrophysiology

#### Recording

Extracellular recordings from layer 5 of barrel columns were targeted as follows: Mice were trimmed to 3 whiskers in C row (C1, C2 and C3) and intrinsic optical imaging was performed to locate the area corresponding to the targeted whisker barrel, under isoflurane anaesthesia (Bray et al., 2016). Acute implantation of a 16-channel linear silicon probe (Cambridge NeuroTech, Cambridge, United Kingdom) was performed at the beginning of each session under isoflurane anaesthesia. Mice were left to recover for at least 2 hours before starting the recording (sampling rate 24.4 kHz, filtering 300-3000 Hz). At the end of a recording session, the microelectrode array was withdrawn, the craniotomy sealed with silicone elastomer, and the mouse returned to its home cage.

### Whisker tracking

After calibration of the cameras, whiskers were tracked in 3D from the video (Petersen et al., 2020). Calibration was performed at the end of each recording session.

The shape of each target whisker was described as a quadratic, 3D Bezier curve ***b***(*s*) = (*x*(*s*), *y*(*s*), *z*(*s*)), defined by 3 control points, where, *x, y, z* are space coordinates and 0 ≤ *s* ≤ 1 parameterises location along the curve: ***b***(*s* = 0) corresponds to the control point closest to the whisker base and ***b***(*s* = 1) to that furthest from the base.

The intrinsic shape of a quadratic curve is fully described by a curvature function *κ*_3*D*_(*s*) (Marsh, 2006):

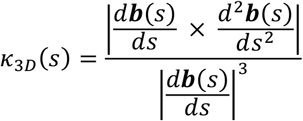

Here |*a* | denotes the magnitude (2-norm) of vector *a. κ*_3*D*_(*s*) has units of 1/distance and is the reciprocal of the radius of the circle that best fits the curve at point *s*. For a quadratic curve, in the quasi-static framework, we can apply the standard relationship between bending moment during contacts (*M*_*z*_) and a change in whisker curvature (Birdwell et al., 2007; Quist & Hartmann, 2012). From here it follows that *M*_*z*_ is proportional to:

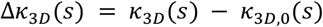

where *κ*_3*D*_(*s*) is the curvature of the whisker during contact and *κ*_3*D*,0_(*s*) is the curvature when the whisker is free from contact and in its resting state. All results presented here were evaluated at *s* = 0 and 5 ms after touch onset.

Bezier curves were visually checked and occasional errors were manually corrected. Touches were manually detected as in Petersen et al (2020).

### Electrophysiological data analysis

#### Spike sorting

Single units (N=33) were isolated from the silicon probe recordings using JRCLUST (Jun et al., 2017). A putative single unit was selected for further analysis if 1) its activity could be tracked during the whole recording, 2) its inter-spike interval histogram exhibited a clear absolute refractory period and 3) its waveform was biphasic and consistent across channels (Campagner et al., 2016).

#### ROC analysis

ROC analysis was carried out as in (Safaai et al., 2013). Responses were measured as spike counts in 30 ms non-overlapping time windows. The hit rate (correct detection of touch) was calculated using response windows starting at touch onset. For first touch, the false positive rate was calculated using ‘noise’ responses sampled between trial onset and first touch. For later touches, ‘noise’ responses were sampled from inter-touch intervals.

#### Tuning curves

To measure tuning curves for Δ*k*_3*D*_, we first identified touch onset times and categorised them as belong to first touch on a trial, second touch, etc. We calculated the mean of Δ*k*_3*D*_ in the 5 ms interval following each touch onset and discarded those with 5% with greatest absolute value as outliers. The remaining samples were discretised into 4 equipopulated bins and, for each unit, the mean spike count (calculated in the 30 ms bin starting at touch onset) for each bin was calculated separately for first touch events, second touch events, etc. For each of these, the tuning curve was estimated as the linear correlation between spike count and Δ*k*_3*D*_.

#### Testing for Sensory Adaptation

When a mouse touches an object by active touch, it controls the force of whisker-object contact and the strength of the touches vary (Results, Figure 2). A change in neuronal response to touch during the course of a sequence of touches might therefore reflect not only SA but also a change in the strength of touch. To tease these effects apart, we first measured the strength of each touch in the trial (first, second etc) as |Δ*k*_3*D*_| and measured the tuning curves separately for each touch. These tuning curves enabled us to estimate (predict) the response of a unit in the state of second touch, for example, to touches of varying |Δ*k*_3*D*_|. To test for SA, we compared the response of units to first touch with that predicted by applying touch strengths measured for first touch through the tuning curve estimated for later touches. This procedure was done using a leave-one-out cross-validation process.

#### How much of the response attenuation during a touch sequence is due to Sensory Adaptation?

The response attenuation evident in Figure 2 potentially reflects both a direct effect of neurons becoming less sensitive (SA) and an indirect effect of motor adaptation weakening the afferent drive. We estimated the contribution of SA in the following way. For robustness, this analysis was carried out using the summed activity of each simultaneously recorded population (‘population response’). For each population, we first calculated the median response to first touch (*FR*_1_) and the median response to later touch (*FR*_*Later*_). The difference *FR*_1_-*FR*_*later*_ is the response attenuation of Figure 2. Next, we used the tuning curve approach described above to ask how much of this attenuation could be attributed to SA alone. To this end, we calculated the tuning curve of the population response to |Δ*k*_3*D*_| for later touches. We then used this tuning curve to estimate the median population response to first touch (*FR*_1_’). In the complete absence of SA, the tuning curve for later touch matches that for first touch, so that *FR*_1_’ is equal to *FR*_1_ and *α* in the following equation is zero:

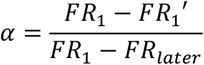

At the other extreme, if the response attenuation effect is entirely due to SA, *α* will equal 1. This index thus quantifies the degree to which response attenuation in a given population can be explained by SA.

#### Stimulus-Specific Adaptation

For this analysis, trials that met all the following criteria were selected: at least the first two touches (the ‘adapting’) were produced by the same whisker (C1 or C2); at least one subsequent touch (the ‘test’) was produced by the other whisker; the time of the test touch did not overlap with that of any other whisker. For comparison between test touch and later (*N*_*touch*_>=3 touch responses, we determined tuning curves for later touches, one for each whisker. These tuning curves were obtained as detailed above, using data for all later touches, except (as a cross-validation procedure) the test stimuli. To quantify changes in the responses to the adapting and test stimuli, we used the index defined in Results (Eq. 1).

